# A Gene Set Foundation Model Pre-Trained on a Massive Collection of Diverse Gene Sets

**DOI:** 10.1101/2025.05.30.657124

**Authors:** Daniel J. B. Clarke, Giacomo B. Marino, Avi Ma’ayan

## Abstract

Trained with large datasets, foundation models can capture complex patterns within these datasets to create embeddings that can be used for a variety of useful applications. Here we created a gene set foundation model that was trained on a massive collection of unlabeled gene sets from two databases: Rummagene and RummaGEO. Rummagene automatically extracts gene sets from supplemental tables of publications; and RummaGEO has gene sets automatically computed from comparing groups of samples from RNA-seq studies deposited into the gene expression omnibus. Several foundation model architectures and data sources for training were benchmarked in the task of predicting gene function. Such predictions were also compared to other state-of-the-art gene function prediction methods and models. One of the GSFM architectures achieves superior performance compared to all other methods and models. This model was used to systematically predict gene functions for all human gene. These predictions are served on gene pages that are accessible from https://gsfm.maayanlab.cloud.

## INTRODUCTION

Foundation models have proven effective in a variety of domains, especially in the natural language processing (NLP) space [1]. The ability of foundation models to disambiguate redundant tokens, encoding them into structured embedding vectors, is their most critical feature. In other words, in NLP, different words or letters are reused in different contexts. Text embeddings have proven useful for many tasks, for example, for semantic text similarity and clustering [2]. The challenges in NLP are like those in biological molecular networks because genes and their protein products are known to play multiple roles, participating in different pathways and biological processes. For example, the gene embeddings created by the Gene2Vec model [3] were found to be useful for predicting gene-gene functional associations when trained on 964 gene-gene co-expression matrices created from the gene expression omnibus (GEO) [4].

More recently, single cell transcriptomics data has been successfully used to build several gene-centric foundation models. For instance, Geneformer [5] is a transformer-based model that was pre-trained with scRNA-seq data collected from ∼30 million single cells, and fine-tuned to perform various tasks such as target identification. Similarly, the foundation model scGPT [6], also a transformer model pre-trained with scRNA-seq data, was demonstrated to be useful for multiple tasks such as cell type annotation, multi-omics integration, perturbation response predictions, and gene-gene network inference. Since scRNA-seq data is noisy and suffers from dropout, Eraslan et al. [7] implemented a Denoising Autoencoder to improve the quality, remove technical variation, and impute missing values of single cell transcriptomics data. These and other similar implementations showed that pre-trained gene-centric foundation models could be fine-tuned to accomplish several useful downstream tasks. However, most of these foundation models are trained with single cell transcriptomics data, covering an important segment of the space of gene modules while potentially missing other important contexts, for example, proteomics, phsphoproteomics, and genomics.

The embeddings of genes into a dimensionality-reduced vector space can be done with various sources of data, for example, protein sequences, gene-gene co-expression correlations, and literature co-mentions. Such embeddings can be used to perform a variety of tasks such as predicting protein-protein interactions, gene-disease associations, and pathway membership [8]. One of the top performing gene embedding models is GenePT [9]. GenePT was created by embedding the National Center for Biotechnology Information (NCBI) gene summaries using an OpenAI top performing large language model (LLM). Another model called Geneshot [10] uses gene-gene co-expression, literature co-mentions, and gene sets submitted to Enrichr [11] to make gene function predictions by computing the average distance of an annotated gene set to all other genes not belonging to the set. The closest genes to the annotated set are predicted to belong to the set. When evaluating the different gene-gene similarity matrices for their ability to make gene function predictions by Geneshot, the annotated sets used for the predictions are divided into two groups: manually curated and omics data derived. As expected, literature-based co-mentions gene-gene matrices perform well in predicting Gene Ontology terms [12] and KEGG pathways [13], whereas co-expression matrices created from transcriptomics and proteomics are better at predicting transcription factor targets [14] and kinase substrates [15].

To improve the quality of the Geneshot predictions, PrismEXP [16] was developed to utilize context specific gene-gene co-expression matrices derived by first clustering samples from ARCHS4, a resource that serves massive collection of uniformly aligned RNA-seq transcriptomics [17]. The collection of these co-expression matrices is used to construct features for a regression model that carries out the gene function prediction task. Benchmarking demonstrated that PrismEXP outperforms Geneshot which in essence uses a single gene-gene co-expression matrix instead of many. The improvement in performance by PrismEXP can be explained by the concept that each ARCHS4 cluster provides context-specific gene-gene association information. While efforts such as ARCHS4 [17] and CELLxGENE [18] provide a rich source for gene embeddings from unbiased transcriptomics experiments across many contexts and conditions, there are many other contexts relevant to gene-gene associations and gene functional space that are missed. For example, results from proteomics, genomics, epigenomics, and many other new and older assays that measure gene and protein expression directly, or indirectly, can provide even greater coverage of gene functional embeddings.

To systematically obtain normalized gene and protein knowledge from diverse resources, Rummagene extracts gene sets by crawling supplemental materials of research publications [19]. While gene sets extracted from supporting tables of research articles do not contain uniformly aligned gene expression data, the extracted gene sets are expected to contain intrinsically varied information about gene sets derived from the variety of experimental conditions and assays. Meanwhile, by using the uniformly aligned bulk RNA-seq data from ARCHS4, RummaGEO [20] also contains a massive collection of gene sets created by computing differential expression between groups of samples from thousands of studies that deposited their RNA-seq expression data into the gene expression omnibus (GEO) [20]. Together, Rummagene and RummaGEO hold a diverse collection of over one million gene sets. Here we present a gene set foundation model (GSFM) trained with the collection of gene sets from Rummagene and RummaGEO. A denoising autoencoder-like model was trained in a self-supervised manner to predict held out genes from unlabeled Rummagene and RummaGEO gene sets. With this model, we show that GSFM outperforms other models and methods with the task of gene function predictions. We make the GSFM model and its predictions available via a web server application and explore other ways the pre-trained GSFM model can be applied for other tasks such as enrichment analysis and protein-protein interactions predictions.

## METHODS

### Data Resources

Various GSFMs were trained with gene sets downloaded from Rummagene [19] and RummaGEO [20] separately and jointly (Table 1). To benchmark the performance of these foundation models, gene-gene similarity matrices were assembled from various sources (Table 2). The names, descriptions, references, and dates when each data resource was accessed are outlined. The final selected GSFM was used to make predictions based on annotated gene sets from several representative gene sets libraries from Enrichr [11] (Table 3).

**Table 1.**
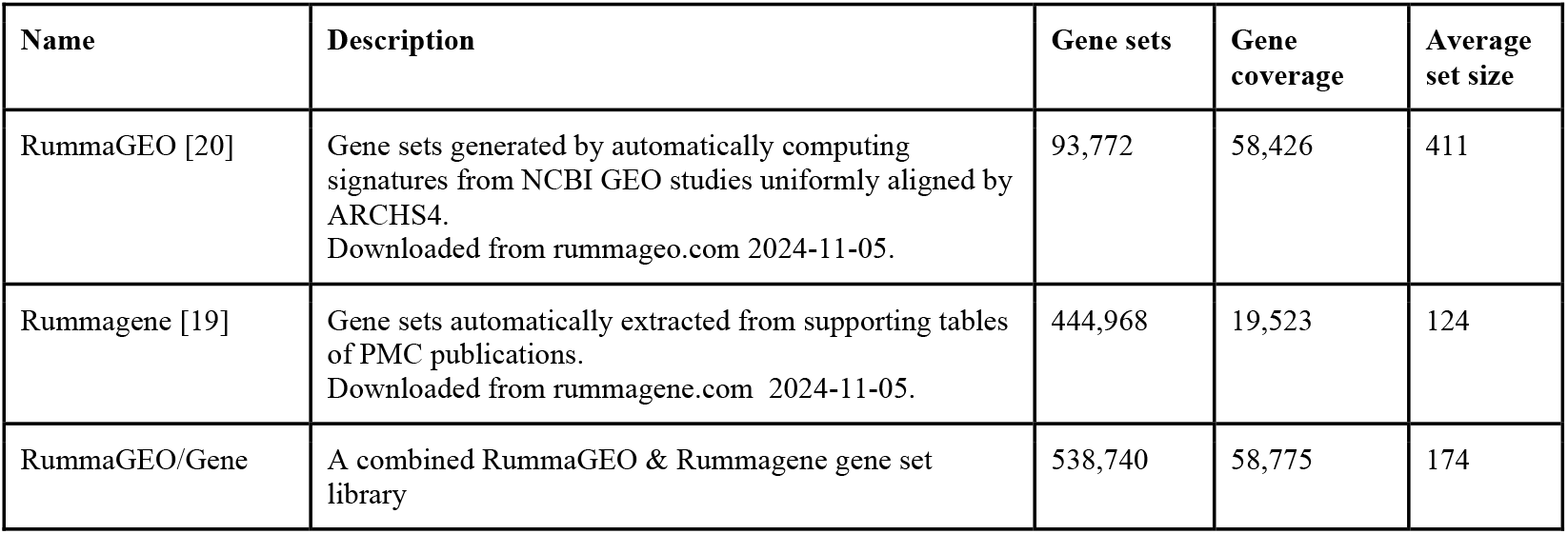
Gene set databases used for training.

**Table 2.** Gene-gene similarity matrices used for gene function predictions.

| Name | Description | Source | Gene coverage |
| --- | --- | --- | --- |
| *ARCHS4* (v2.4) [17] | Gene-gene co-expression correlations computed with Pearson correlation coefficient (PCC) gene across uniformly aligned samples. Downloaded from ARCHS4 2024-10-08. | 855,835 RNA-seq GEO samples | 29,082 |
| *GTEX* Tissue Gene Expression Profiles [24] | Cosine similarity of tissue expression associations for pairs of genes downloaded from Harmonizome [25] 2024-10-08. | 15,201 RNA-seq samples collected from 49 tissues of 838 donors | 17,369 |
| *HPA* Tissue Protein Expression Profiles [25,26] | Cosine similarity of immunohistochemistry protein expression associations for pairs of proteins from HPA. Downloaded from Harmonizome [25] 2024-10-08. | 44 normal human tissues, samples from 144 individuals (at least 3 individuals for each tissue) | 15,706 |
| *Enrichr* Gene-Gene Co Occurrence from Submitted Sets [11] | Gene co-occurrence from user submitted gene lists were downloaded from Geneshot [10] 2024-11-04. Co-occurrence counts were normalized by the diagonal. | 293,747 enrichr user-submitted gene sets | 23,568 |
| *GeneRIF* Gene-Gene Co Occurrence [27] | Gene co-occurrence from NCBI's GeneRIF were downloaded from Geneshot [10] 2024-11-04. Gene-PubMed ID associations from GeneRIF were downloaded from the NCBI's GeneRIF FTP server. | 1,015,165 gene-publication pairs for 647,803 publications | 16,729 |
| *Tagger* [28] | Gene co-occurrence in PubMed abstracts. Co-occurrence counts were normalized by the diagonal. Downloaded from Geneshot 2024-11-04. | 9,353,632 gene-publication pairs | 20,952 |
| *GenePT* [9] | Gene-gene similarity based on embeddings produced with OpenAI applied to NCBI gene definitions. Gene-gene similarity is computed with cosine similarity. Download from figshare 2025-02-10. | 29,875 gene embeddings | 29,875 |
| *RummaGEO* [20] | Gene sets generated by automatically computing signatures from NCBI GEO studies uniformly aligned by ARCHS4. | 50,000 random gene sets | 19,484 |
| *Rummagene* [19] | Gene sets automatically extracted from supporting tables of PMC publications. | 50,000 random gene sets | 19,528 |
| *RummaGEO & Rummagene* | Computed by cosine similarity of 50k random gene sets from RummaGEO multi-hot encoded and 50k random gene sets from Rummagene multi-hot encoded. | 100k random gene sets | 60,660 |

**Table 3.**
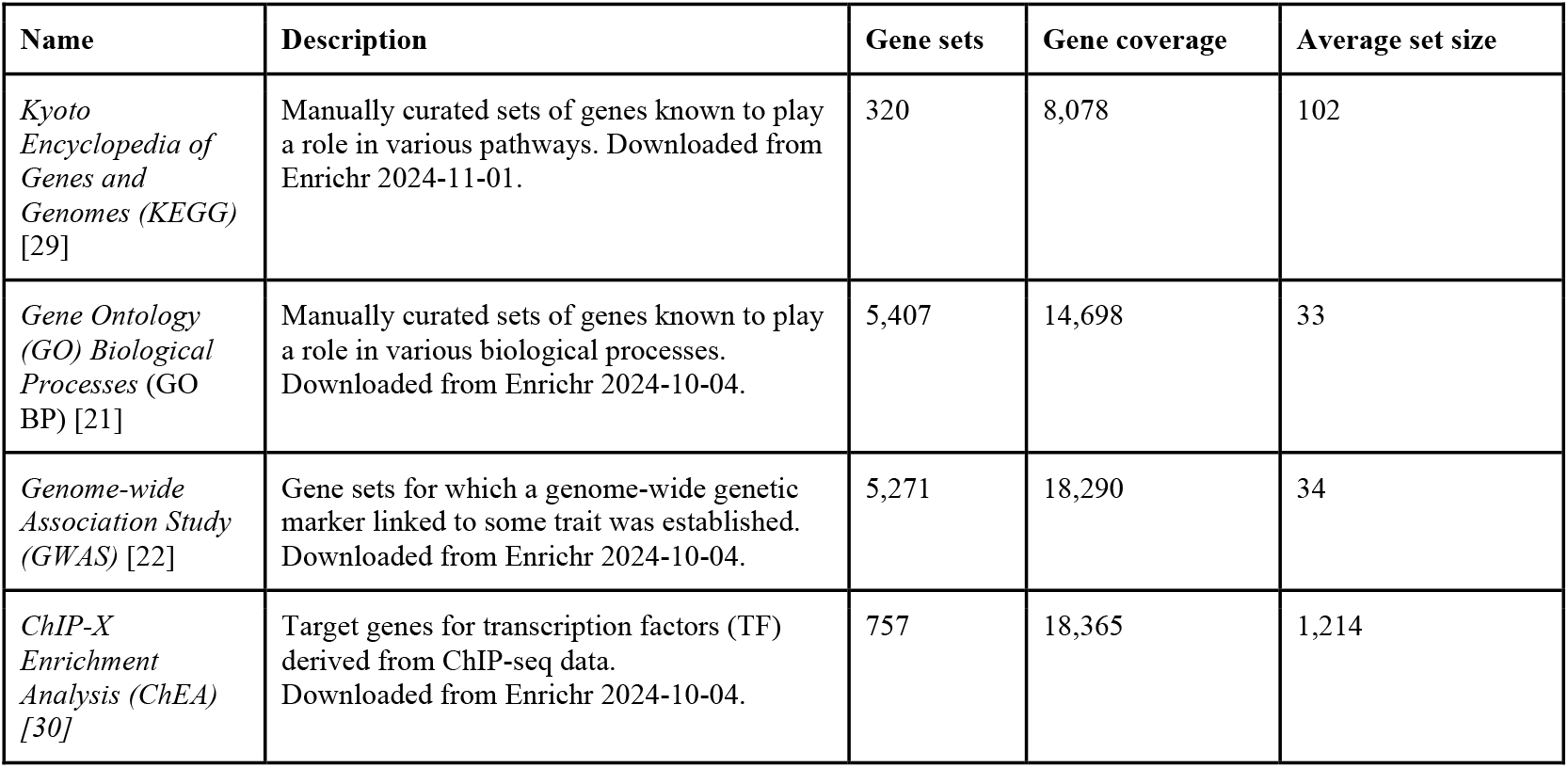
Gene set libraries used for benchmarking.

| Name | Description | Gene sets | Gene coverage | Average set size |
| --- | --- | --- | --- | --- |
| *Kyoto Encyclopedia of Genes and Genomes (KEGG)* [29] | Manually curated sets of genes known to play a role in various pathways. Downloaded from Enrichr 2024-11-01. | 320 | 8,078 | 102 |
| *Gene Ontology (GO) Biological Processes* (GO BP) [21] | Manually curated sets of genes known to play a role in various biological processes. Downloaded from Enrichr 2024-10-04. | 5,407 | 14,698 | 33 |
| *Genome-wide Association Study (GWAS)* [22] | Gene sets for which a genome-wide genetic marker linked to some trait was established. Downloaded from Enrichr 2024-10-04. | 5,271 | 18,290 | 34 |
| *ChIP-X Enrichment Analysis (ChEA)* [30] | Target genes for transcription factors (TF) derived from ChIP-seq data. Downloaded from Enrichr 2024-10-04. | 757 | 18,365 | 1,214 |

### Absolute Average Similarity for Gene Function Prediction

Using the benchmarking gene-gene similarity matrices (Table 2), gene function predictions were performed using a simple correlation-based method as previously described for the Geneshot resource [10]. Briefly, all genes that are not in the annotated gene set are assigned a score based on their normalized average absolute correlation with the genes already in each annotated gene set.

### PrismEXP Gene Function Prediction

The PrismEXP Python package [16] was used to make gene function predictions with various background datasets. PrismEXP first partitions a data matrix where genes are the rows and samples are the columns into clusters. It then creates a Pearson gene-gene similarity matrix for each cluster. In the second step, the PrismEXP algorithm uses a gene set library to create features for a classifier by converting each set in the library to all-gene vectors based on the average distance of all genes to the genes in each set within the gene set library. This process is repeated for each cluster. The original implementation of the PrismEXP algorithm used the uniformly aligned bulk RNA-seq data from ARCHS4 [17] and the Gene Ontology Biological Processes (GO BP) [21] gene set library created for Enrichr [11]. This approach and implementation are included as a benchmark here. To compare the PrismEXP method with other methods applied to the same task of gene function prediction more broadly, including GSFM, we also applied the PrismEXP algorithm to a one-hot-encoded gene set matrix created from Rummagene [19] for the first step of the PrismEXP algorithm. For the second step, we used a sampled Rummagene gene set library made of 10,000 gene sets randomly selected from all 100 clusters identified in the first step.

### GSFM Architectures

Several denoising autoencoder-like model architectures (Fig. 1) were pre-trained in a self-supervised fashion. During training, the models optimized a binary cross-entropy loss function by predicting a set of genes as a multi-hot encoded vector. Below we describe the various architectures that were trained and benchmarked:

**Figure 1.**
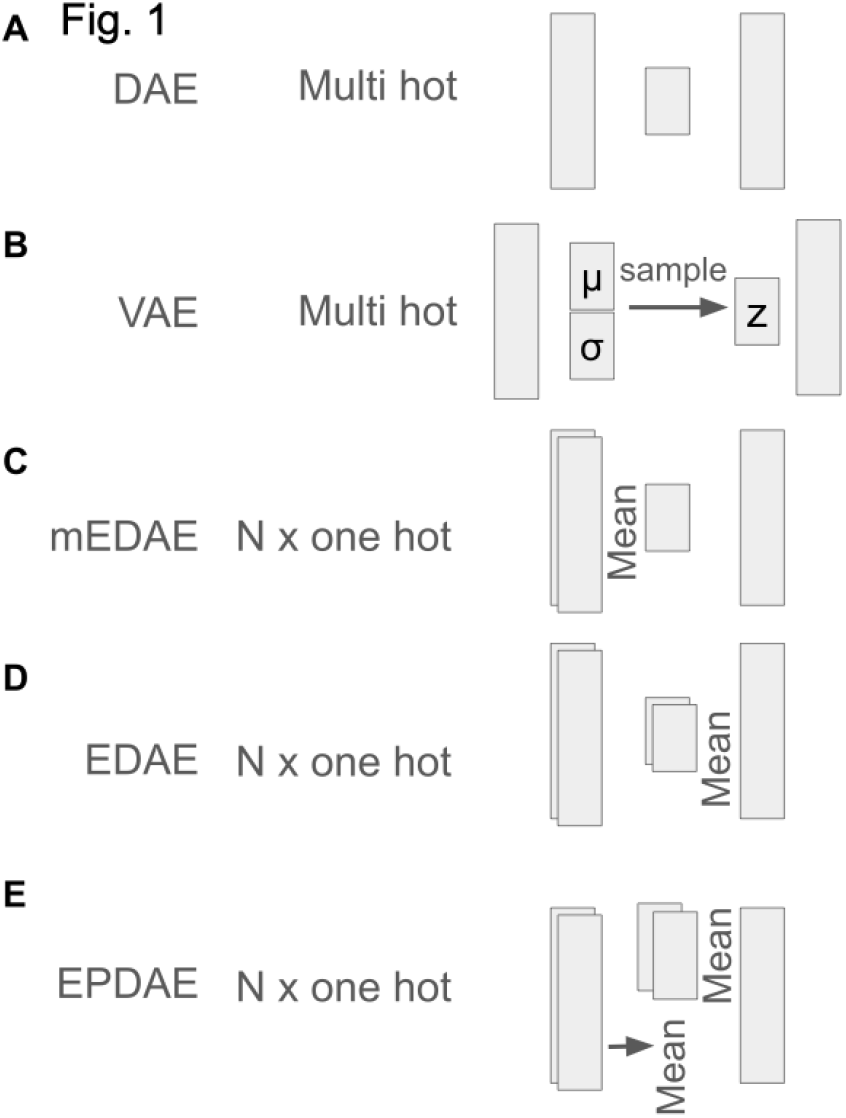
GSFM Architectures. The denoising autoencoder-like architecture variations illustrate how each variant collapses the network to a single vector at some place prior to the final prediction.

A standard Denoising Auto Encoder (DAE) (Fig. 1A) is trained with the multi-hot-encoded input gene set. The symmetric network consists of a sequence of progressively smaller layers by a factor of two until the bottleneck layer. For the bottleneck, the decoder acts as a multi-label classifier. The model is trained to minimize the binary cross-entropy loss between the predicted labels and the unseen genes in the set. A standard Variational Autoencoder (VAE) (Fig. 1B) was also trained with the same binary cross-entropy reconstruction loss but with the added KL divergence term.

Several variations of DAE were also trained, tested, and benchmarked. These variations use a linear embedding layer instead of a multi-hot encoded vector as the input layer. Since this results in a vector for each gene in the set instead of a single vector, several architectures were designed and implemented based on how these models collapse the multiple vectors into one. For the Mean Embedding Denoising Auto Encoder (mEDAE) architecture (Fig. 1C), the embeddings are averaged before the encoder. For the Embedding Denoising Auto Encoder (EDAE) model (Fig. 1D), the mean is applied at the end of the encoder. For the Embedding Passthrough Denoising Auto Encoder (EPDAE) architecture (Fig. 1E), both the mean embedding and the mean after the encoder are concatenated to produce the latent layer feeding into the decoder.

All these models were tested and hyper-parameterized manually. The hyperparameters include 1) dropout: applied to all layers during training; 2) partitioning: the amount that was hidden to complete the gene set; 3) d_model: the size of the latent space and embedding layers; 4) depth: the number of hidden layers between the input and the latent space; and 4) weighted_loss: whether or not the binary cross-entropy loss function weights the loss by the inverse of that gene’s prevalence in the dataset. Each model and set of hyperparameters were trained and evaluated for at least 20 epochs. Checkpoints were created every 5 epochs during training. Models were trained with gene sets from Rummagene [19] alone, from RummaGEO [20] alone, and with all the sets from both Rummagene and RummaGEO, and these combinations were evaluated on the benchmarking libraries.

### Benchmarking Models and Datasets

All gene function prediction methods including the absolute average similarity, PrismEXP, and the GSFM models were used to predict other genes that should belong to a set, given an annotated gene set. For each gene set in each benchmarking library (Table 3), gene sets were randomly shuffled, and each gene set was split in half five times. Conditioned on one half, the methods were used to predict the missing genes for each gene set. The methods were evaluated by their ability to recover the missing genes from the hidden other half. The area under the receiver operating characteristic (AUROC) curve is computed for each method and each benchmarking library. The median AUROC across the random splits for each term are computed, and the distribution of median AUROCs across the entire library for each method are visualized side-by-side. To account for the different vocabulary sizes between methods, only predictions for genes already present in each gene set library were considered when evaluating the different methods.

## RESULTS

### Baseline Predictive Performance

To establish a baseline for predictive performance of different models and training datasets, the absolute average similarity approach for gene function prediction [11] was implemented to illustrate the performance of different training data sources (Fig. 2). The GenePT [9] similarity matrix shows the best performance in predicting missing genes from manually curated gene set libraries, namely GO BP [12] and KEGG [13]. On the other hand, prioritizing missing genes for gene sets created from data driven gene set libraries, namely GWAS Catalog [22] and ChEA [14], is best achieved with gene-gene similarity matrices created from RummaGEO [20], Rummagene [19], and Enrichr [11,19].

**Figure 2.**
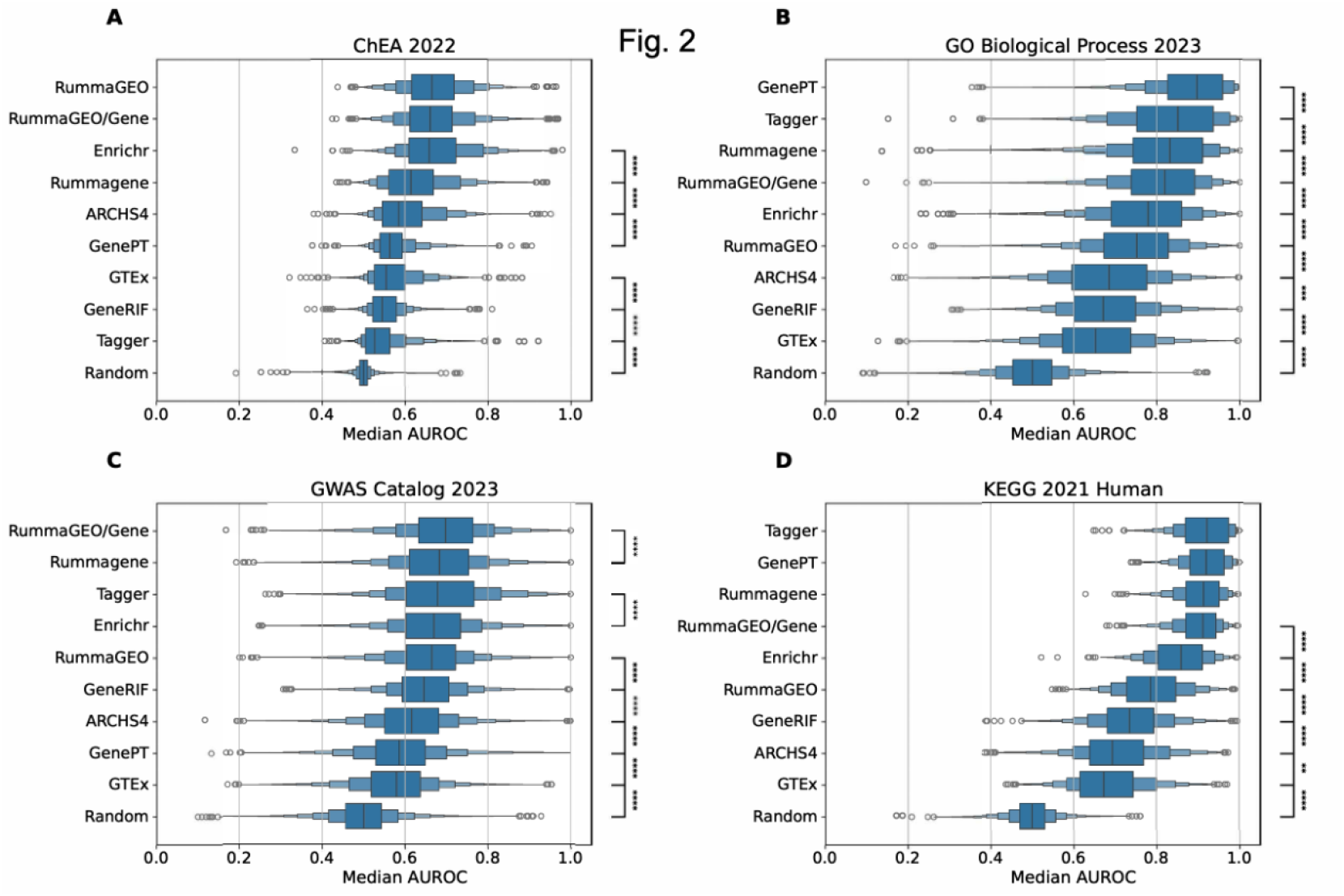
Baseline Benchmarks. Performance of absolute average similarity applied on gene-gene similarity matrices derived from different data resources and modalities. Sorted by average performance, significance testing performed between consecutive distributions.

### Benchmarking GSFM Model Architectures

The various GSFM model architectures were trained with the Rummagene and RummaGEO gene sets (Table 1) using different hyper parameters for 50 epochs and evaluated for their performance. The partitioning hyperparameter determines the portion of genes from each gene set that is made visible to the model. For example, a value of 0.2 would mean that 80% of the genes in the set are given to the model for predicting the remaining 20%. The special value of 0 means that the model is provided with all genes in each set. We found mixed results for different partitioning schemes (Fig. 3A). For dropout, or in other words for the random zeroing of weights throughout the network during training, different values have different effects on the different benchmarks with the value 0.2 performing well overall (Fig. 3B). The network depth corresponds to the number of layers from the input to the latent layer. A depth of 2, which is only one hidden layer, seems to perform best overall (Fig. 3C). A dimensionality corresponding to a hidden layer of 256 dimensions seems to work best across benchmarks (Fig. 3D). Re-balancing the loss function based on the frequency of genes, seems to have negative effects on performance (Fig. 3E). Although several other model architectures were tested, the standard multi-hot DAE seems to outperform all alternative approaches across the four benchmarking libraries (Fig. 3F). Given these hyper-parameter selections for the final selected GSFM, this model was trained for 100 epochs. We found that the predictive quality of this GSFM levels off at around 50 epochs (Fig. 3G). We then trained the same GSFM architecture with gene sets from RummaGEO [20] and with a mixture of both, finding that training the GSFM with only the Rummagene [19] gene sets broadly outperforms training the model with additional and alternative gene sets from RummaGEO (Fig. 3H).

**Figure 3.**
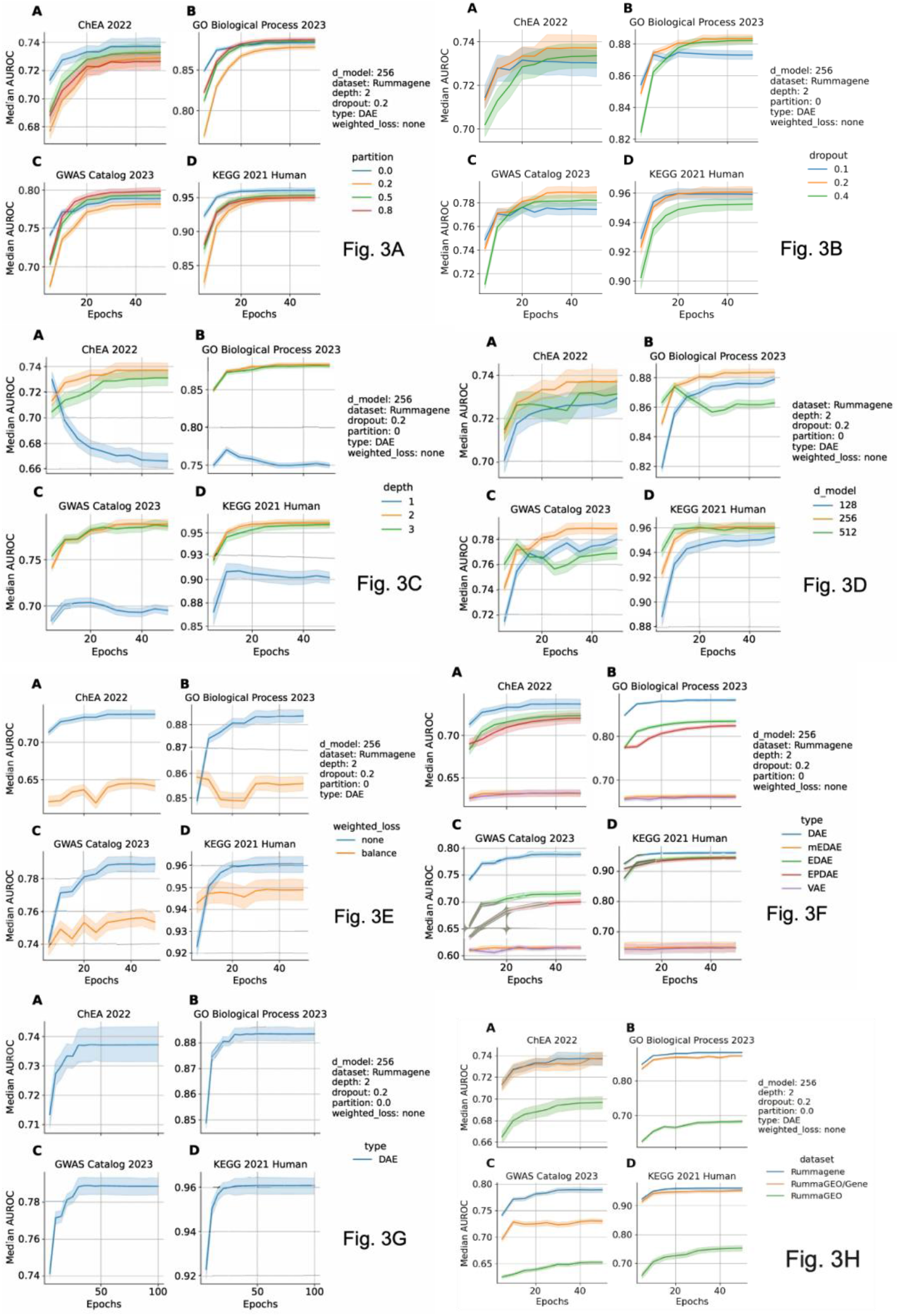
Evaluation of Different Architectures, Datasets, and Hyperparameters. A) Evaluating partitioning thresholds for training and testing gene set completion. B) Evaluating dropout levels. C) Evaluating depths of model layers. D) Evaluating bottleneck vector sizes. E) Testing whether to apply weighted loss balancing. F) Evaluating architectures. G) Training the top performing model with more epochs. H) Evaluating top performing model with different training datasets.

### Predictive Performance by Method and Application to Other Libraries

To benchmark the final selected GSFM with other models and methods that can be used to predict gene function, the AUROC curves across all methodologies and datasets are compared (Fig. 4). We see that the more complex algorithms such as PrismEXP [16] and GSFM perform better than the absolute average similarity when using the same underlying data. For example, across all benchmarks, GSFM trained with Rummagene (“GSFM Rummagene”) does better than PrismEXP parameterized by Rummagene (“PrismEXP Rummagene/Rummagene”), which does better than absolute average similarity of the Rummagene’s gene-gene similarity matrix (“Rummagene”). However, GSFM trained with Rummagene outperforms all other tested methodologies and datasets across the four benchmarking libraries. Since GSFM appears to have superior performance, we applied the GSFM, trained with Rummagene, to make predictions with a wide variety of additional libraries (Table 4). Adopting the same methodology used for benchmarking, we evaluated the median AUROC curves across these additional libraries (Fig. 5). Then, for each gene set from these additional gene set libraries predictions were made for all human protein coding genes giving to the model all known genes for each set. These predictions are made available in gene-centric landing pages. These landing pages also include a textual summary about each human gene from NCBI. Alternatively, by giving an LLM (Gemini or GPT-4o) the top 50 most cited articles that mention the gene name in their abstracts, gene descriptions with citations are also provided on the gene landing pages.

**Table 4.** Gene set libraries used for GSFM predictions.

| Name | Description | Gene sets | Gene Coverage | Average Set Size |
| --- | --- | --- | --- | --- |
| *KOMP2 Mouse Phenotypes 2022*<br>*10.1093/nar/gkac972* | Downloaded from CFDE Workbench 2025-03-03 | 529 | 6,713 | 68 |
| *MoTrPAC Endurance Trained Rats 2023*<br>*10.1016/j.cell.2020.06.004* | Downloaded from CFDE Workbench 2025-03-03 | 225 | 10,428 | 115 |
| *KEA 2015*<br>*10.1093/nar/gkab359* | Downloaded from Enrichr 2025-03-03 | 428 | 3,102 | 25 |
| *LINCS L1000 Chem Pert Consensus Sigs* | Downloaded from CFDE Workbench 2025-03-03 | 10,850 | 10,850 | 245 |
| *MGI Mammalian Phenotype Level 4 2024* | Downloaded from Enrichr 2025-03-03 | 5,003 | 11,625 | 36 |
| *LINCS L1000 CRISPR KO Consensus Sigs*<br>*10.1093/nar/gkac328* | Downloaded from CFDE Workbench 2025-03-03 | 10,424 | 9,440 | 245 |
| *LINCS L1000 Consensus Median Signatures*<br>*10.1093/nar/gkac328* | Downloaded from CFDE Workbench 2025-03-03 | 16,936 | 9,792 | 200 |
| *Human Phenotype Ontology*<br>*10.1093/nar/gkaa1043* | Downloaded from Enrichr 2025-03-03 | 1,779 | 3,096 | 31 |
| *HuBMAP Azimuth* | Downloaded from Enrichr 2025-03-03 | 1,425 | 3,712 | 9 |
| *HuBMAP ASCT+B* | Downloaded from CFDE Workbench 2025-03-03 | 351 | 874 | 8 |
| *OMIM Disease* | Downloaded from Enrichr 2025-03-03 | 90 | 1,759 | 25 |
| *IDG Drug Targets 2022* | Downloaded from CFDE Workbench 2025-03-03 | 888 | 1,552 | 16 |

**Figure 4.**
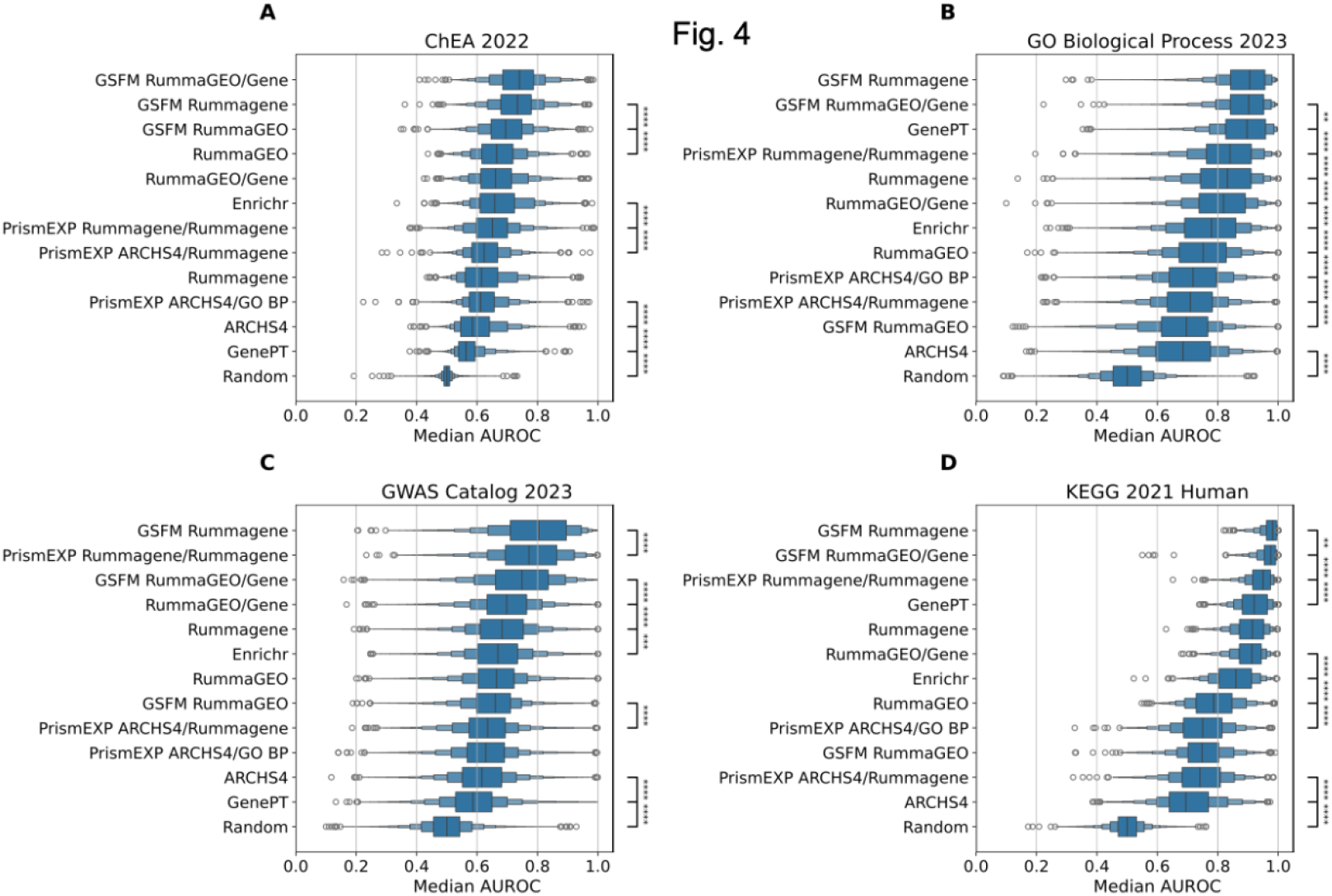
Performance of Methods based on Data Derived from Rummagene and RummaGEO. Sorted by average performance, significance testing performed between consecutive distributions.

**Figure 5.**
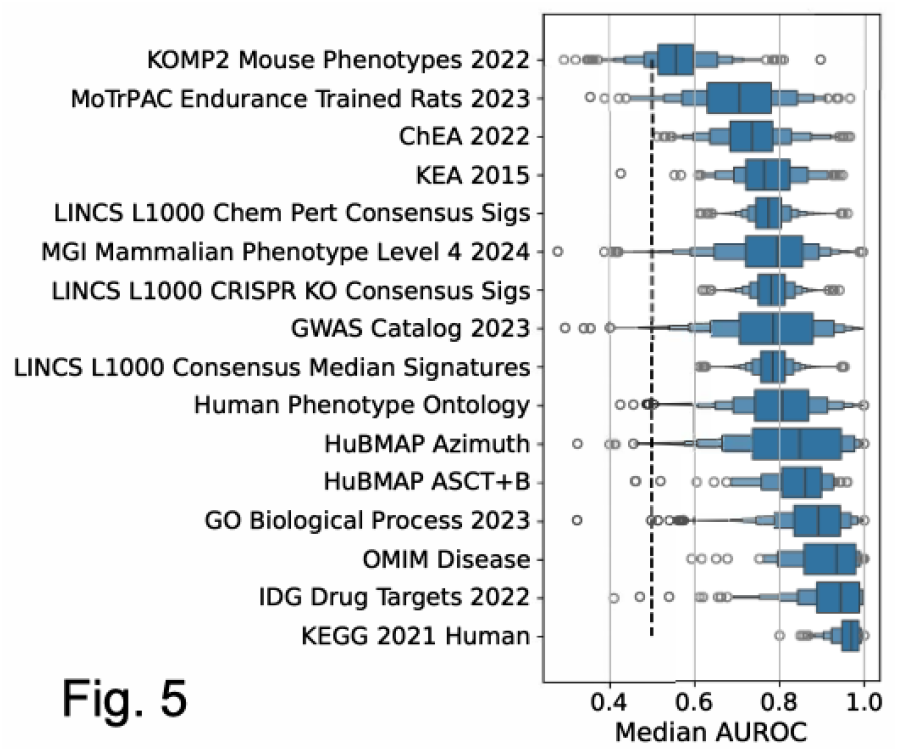
Evaluation of Predictions for Additional. Performance distribution of Rummagene-trained GSFM on libraries used to make predictions for the GSFM website.

### Applying the Model to Downstream Tasks and Making it Available

To make GSFM more accessible to the community, we developed a user-friendly website that hosts the model, the processed datasets and gene sets, and the predictions made by the model. The site is available from: https://gsfm.maayanlab.cloud/ and it enables users to browse pre-computed predictions for all human genes across multiple libraries (Fig. 6). Model-assigned logits, the logit z score, and information for calibrating expectations of generalizability were included in the form of AUROC curves for each gene set library term, the number of genes that were predicted to be associated with that term, and the number of genes where that term would appear in the top 10 predicted terms. The GSFM site also enables users to perform zero-shot gene function prediction with the pretrained GSFM model, as well as the ability to augment any gene set with additional genes using the model. The pre-trained model is also published on HuggingFace at: https://huggingface.co/maayanlab/gsfm for the community to apply for other downstream tasks. GSFM should prove useful for gene function prediction, gene set embedding, and as a pre-trained embedding for training other models that include gene sets.

**Figure 6.**
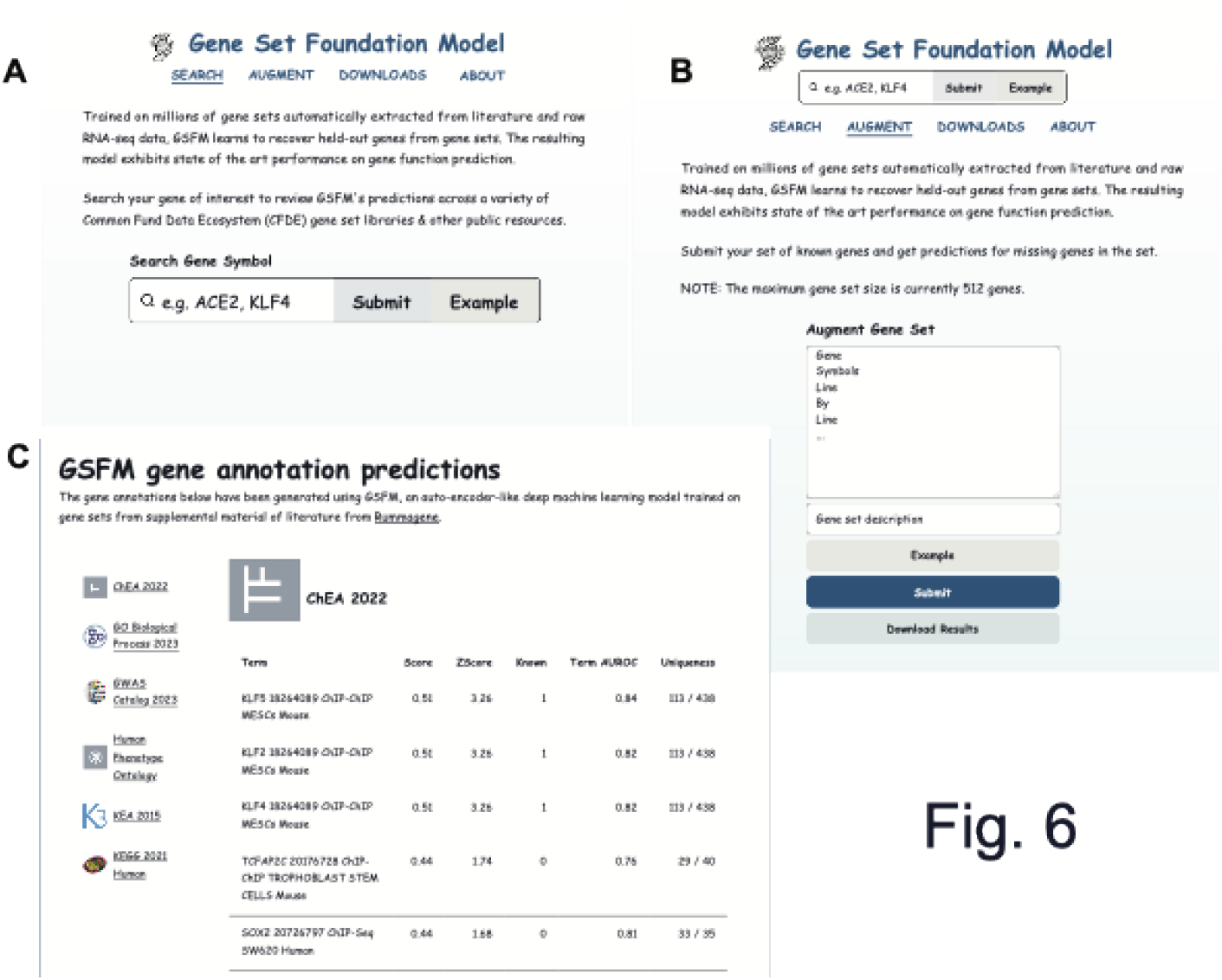
Screenshots from the GSFM Website. (A) the home page, (B) the augment page (C) the gene pages, and (D) the predictions on the gene pages.

## DISCUSSION

GSFM’s ability to more accurately predict genes held out from known gene sets can be useful for many applications in computational systems biology. Unlike prior methods that are mainly based on similarity of all genes to annotated genes, GSFM’s architecture can capture the more complex non-linear and multimodal relationships between genes and the gene modules these genes constitute. We demonstrate that various foundation model architectures trained on many unlabeled gene sets can make better gene function predictions when compared with existing methods. We also show that the diversity of gene sets collected for the Rummagene resource [19] can be leveraged to make high quality predictions about gene function based on both data-driven and manually curated sources.

Besides predicting gene function based on annotated gene set completion, GSFM can be used for other tasks, for example, predicting protein-protein interactions, kinase-substrate phosphorylations, annotation of single cell types from expression profiling, and gene set enrichment analysis [23].

Furthermore, GSFM is currently only trained on unlabeled gene sets. However, since there is rich textual information about each gene set from Rummagene [19], RummaGEO [20] and other large-scale sources of gene sets, i.e. Enrichr [11], the next obvious step for GSFM is to develop gene set foundation models that include the gene set annotations as a part of the inputs and outputs of such foundation models. The pre-trained GSFM embeddings could be used as one part of such models. Another limitation of GSFM is that it is trained with only coding genes. Training a similar model using all known genes, coding, and non-coding, as well as different isoforms and transcripts can be useful for many other downstream tasks and applications. GSFM is also trained using mammalian genes, namely human and mouse. However, the same approach can be applied to available gene sets from other model organisms. Finally, many sources of gene and protein sets are in the format of signatures. These signatures can be represented as up/down gene sets, or as vectors of genes with an associated value, for example, differential expression p-values. Training foundation models with such input vectors should be possible and can be useful for applications not currently covered by GSFM. In summary, GSFM’s ability to distill knowledge from large amounts of unlabeled gene sets automatically, and to do so successfully across multiple sources of knowledge, can be translated into many low-hanging-fruit hypotheses that could be tested in wet lab experiments to rapidly advance knowledge in biomedical research.

## RESOURCES AVAILABILITY

The GSFM website, providing access to predictions and the ability to use the model can be found at: https://gsfm.maayanlab.cloud

All benchmark results are available at: https://gsfm.maayanlab.cloud/downloads

The foundation model source code and weights with instructions how to access it locally are available on HuggingFace at: https://huggingface.co/maayanlab/gsfm

The source code for the website is available at: https://github.com/maayanlab/gsfm-predictions

